# ORCO: Ollivier-Ricci Curvature-Omics — an unsupervised method for analyzing robustness in biological systems

**DOI:** 10.1101/2024.10.06.616915

**Authors:** Anish K. Simhal, Corey Weistuch, Kevin Murgas, Daniel Grange, Jiening Zhu, Jung Hun Oh, Rena Elkin, Joseph O. Deasy

## Abstract

Although recent advanced sequencing technologies have improved the resolution of genomic and proteomic data to better characterize molecular phenotypes, efficient computational tools to analyze and interpret the large-scale omic data are still needed. To address this, we have developed a network-based bioinformatic tool called Ollivier-Ricci curvature-omics (ORCO). ORCO incorporates gene interaction information with omic data into a biological network, and computes Ollivier-Ricci curvature (ORC) values for individual interactions. ORC, an edge-based measure, indicates network robustness and captures global gene signaling changes in functional cooperation using a consistent information passing measure, thereby helping identify therapeutic targets and regulatory modules in biological systems. This tool can be applicable to any data that can be represented as a network. ORCO is an open-source Python package and publicly available on GitHub at https://github.com/aksimhal/ORC-Omics.

## Introduction

Recent advanced sequencing technologies enable researchers to investigate biological phenotypes related to a disease through whole genome-wide landscape scans. As a result, there has been unprecedented increase in high resolution biological data such as RNA-seq, DNA-seq, and proteomic sequencing data. A number of tools have been developed in the field of bioinformatics to analyze the large-scale data, including DESeq2, edgeR, GSEA, and UMAP [1,2]. However, bioinformatic tools with the capability that assesses the robustness of interactions via the entire biological systems are still lacking. Biological systems can be represented as undirected graphs where nodes represent a key component and edges represent interactions or correlations between components. Examples include genomic networks, where genes represent nodes and gene interaction information defines edges, and proteomic networks, where proteins represent nodes and expression correlation levels represent edges.

A key goal of analyzing biological systems is understanding critical genes for a given data context as well as the behavior of their local neighborhoods. There are many network analysis methods that can be applied, such as node connectedness, centrality, and eigenvector centrality [3]. However, graph analysis tools with the capability to assesses the robustness of interactions including the entire system are still needed. To address this, we have developed a network-based tool, called Ollivier-Ricci curvature-omics (ORCO). ORCO utilizes Ollivier-Ricci curvature (ORC), an extended notion of Ricci curvature defined on a Riemannian manifold [4]. Given a biological network, ORCO measures edge weights using a consistent information passing measure in neighborhoods of the two nodes. This metric reflects more realistic biological interactions compared to simple correlation between two nodes. Therefore, the network-based approach could provide unique and unexplored insights into the underlying biology and help identify therapeutic targets and regulatory modules within biological systems [5].

Examples of robust features include redundancy and feedback loops. For example, redundancy ensures that an electrical power grid keeps transmitting even if an individual power station is down. In a protein-protein interaction network, the feedback behavior may enable a cell to develop drug resistance. A lack of robust connectivity between network nodes may indicate a vulnerability in coordinated action. In principle, ORC can be considered to quantify the number of pathways between two nodes. The more pathways exist between a pair of nodes, the more robust the relationship between the two nodes is considered to be. Inversely, if removing a single edge in a network makes it impossible for information to pass through, then that edge is considered fragile. Long-range changes in robustness indicate system-level changes in functional cooperation, which may bear important biological implications. For example, abnormalities in robustness may translate to potential therapeutic targets [6].

ORC has been used in various biomedical studies, including brain connectivity analysis and genomic network analysis in cancer cells [7,8]. In [9,10], ORC was used to elucidate differences in brain connectivity of children with autism spectrum disorders (ASD). It was shown that the information provided by ORC was not simply complimentary to the information displayed by alternative methods but uncovered previously unknown connectivity patterns. In oncology, ORC has been used in a variety of applications. In [11], ORC was used to identify a novel gene signature for high-risk multiple myeloma. In [12], a dynamic form of ORC was used to identify therapeutic targets in sarcoma. In [13–15], ORC was used to identify robust genes while taking into account various types of interaction data. ORC also has been shown to improve the performance of neural networks. In [16], the authors used ORC to improve the messaging-passing capabilities of their graph neural networks. ORC was also used for community detection [17]. Together, these works demonstrate that the curvature of biological networks can serve as a prognostic biomarker for various disease states, identify previously unknown high-risk patient groups, and rank their unique geometric vulnerabilities.

Here, we present ORC-Omics (ORCO), an open-source Python library explicitly designed for typical bioinformatics network analysis. ORCO incorporates node data and node adjacency and outputs a network where edge weights represent the robustness between nodes. Additionally, while ORCO is designed for omic data, it can be used in any context where data is represented as a network.

### Software

The following sections provide a technical overview of the method and input data requirements. We describe the input data using genomic data as an example but any appropriate data type could be used.

#### Input data

ORCO requires two input data. The first is a 2D matrix of non-negative feature data where the columns represent samples and rows represent features (e.g., genes). The second is a 2D adjacency matrix where the dimensions match the number of features in the dataset. This adjacency matrix should be a binary symmetric matrix where a positive value indicates an interaction between features. Other input parameters associated with the ORC formulation may be modified and described in more detail on the GitHub page. An example input data structure would be a comma-separated values (CSV) file with *n* columns where each column represents a sample and *m* rows where each row represents a gene. Depending on the data modality, the value of each cell could be gene expression or other omic data.

For genomics data, adjacency matrixes can describe genes that are known to physically interact, co-express, or are connected. For omics data, the matrixes can represent any appropriate type of interaction. Common sources of protein interaction information include the Human Protein Reference Database [18] (HPRD) and the String protein-protein interaction network [19].

#### Ollivier-Ricci Curvature

The following sections describe the both the computation of edge probabilities from node data and the implementation of ORC used.

A notion of graph distance, defined between every two nodes in the network, is required for computing ORC. The standard hop distance may be used. Alternatively, a more informed weighted hop distance can be computed by considering a random walk on the weighted graph as follows. For each sample, the weights *w*_*i*_ for each node *i* are assigned by mapping the genomic data for each gene to the corresponding node in the graph. The node weights are then used to define transition probabilities from one gene to another, which is only non-negative if there is an edge (i.e., protein interaction) between the corresponding genes. The transition probability from gene *i* to any neighboring gene *j*, denoted *p*_*i,j*_, is computed based on the mass action principle normalized over all neighbors to ensure it is a probability, defined as follows:

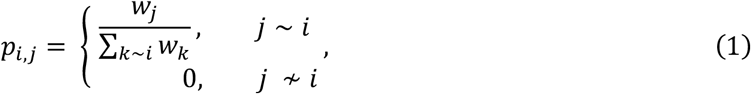

where *k* ∼ *i* denotes the set of nodes *k* that have an edge connecting to node *i*.

Considering that the higher the probability of transitioning from one gene to another, the smaller the edge length should be, the edge weights *w*_*ij*_ for each edge (i, j) are defined as a function of the transition probabilities in each direction as shown in Equation 2:

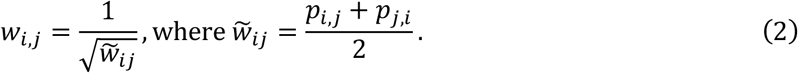

The weighted hop distance between any two nodes *i* and *j*, denoted *d*(*i, j*), is then the minimum accrued weight over all paths connecting nodes *i* and *j*.

Formally, ORC between any two nodes *i* and *j* is defined in Equation 3:

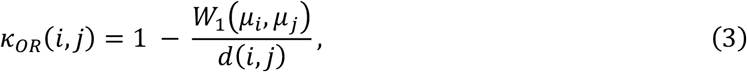

where *W*_1_ is the Wasserstein distance, also known as the Earth Mover’s distance (EMD), between the probability distributions, *μ*_*i*_ and *μ*_*j*_ associated with nodes *i* and *j*, respectively. The probability distribution around a given node (gene), *μ*_*i*_, is defined by the mass action principle, normalized by the net action over all of its neighbors, as follows:

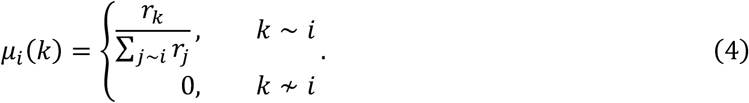

Here *r*_*k*_ denotes the weight of node *k* (e.g., the RNA-Seq value), and the denominator *d(i, j)* is the weighted shortest path between the two nodes, described above.

The output of this method is a graph with an ORC value for each edge. This output can be analyzed in several ways. Figure 1 illustrates an example that shows the cascading effects of overexpression of a single gene throughout the network. ORCO can be installed via Python’s package installer ‘pip.’ The source code, documentation, and examples can be accessed at github.com/aksimhal/ORC-Omics. ORCO represents a powerful tool for researchers to analyze genomic data through a network lens.

**Figure 1:**
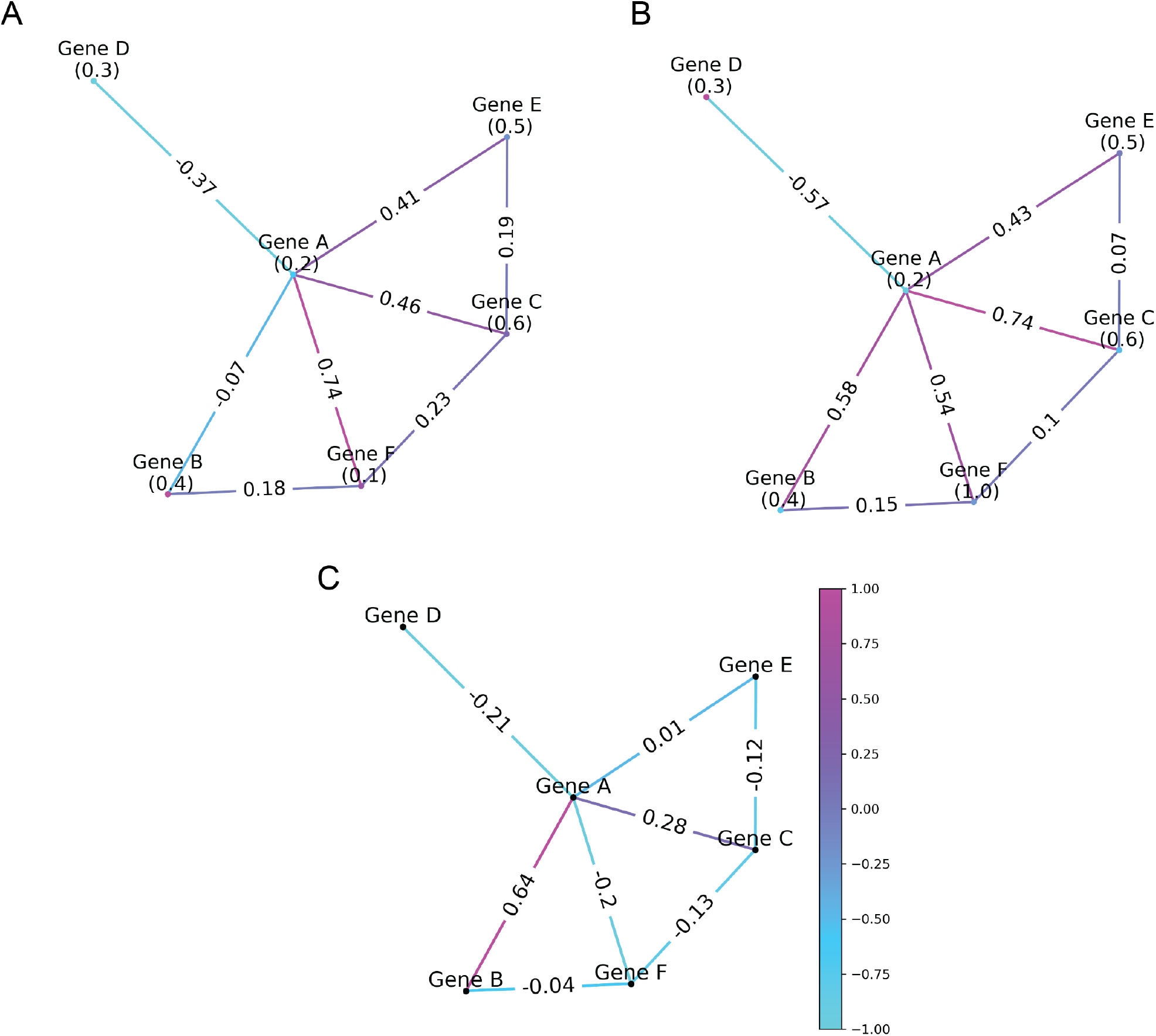
ORCO example. Each subfigure has the same topology and node weights, except for node F. In (A), *Gene F* has a weight of 0.1. In (B), *Gene F* has a weight of 1.0. (C) shows the change in robustness (network B – network A) that cascade throughout the network based on an overexpression of *Gene F*. For example, as *Gene F* expression increases, the connection between *Gene D* and *Gene A* becomes more fragile. On the other hand, the connection between *Gene B* and *Gene A* becomes more robust.

### Runtime considerations

ORC requires the solution of OMT problems between every pair of nodes and hence computation time can be significant. The problem is symmetric in origin/target nodes and can be easily parallelized and ORCO uses multiple cores if provided. Typical runtime for a 500-node, 31,000 edge random Erdős–Rényi graph with random node weights is approximately 12 seconds on an 8-core 8 GB M1 Apple MacBook Pro.

## Conclusion

ORCO is an open-source tool that provides an easy entry point into Wasserstein-based network analysis. As shown in prior publications, ORC has helped discover novel oncological insights; however, the depth of insight ORC can provide is underexplored, and there are still many cancer types for which ORC has yet to be thoroughly investigated. We hope this tool will assist other researchers with their science.

## Acknowledgments

This study was supported in part by an MSK Cancer Center Support grant (P30 CA008748), The Simons Foundation, and a Breast Cancer Research Foundation grant (BCRF-17-193).

